# T-SNE visualization of large-scale neural recordings

**DOI:** 10.1101/087395

**Authors:** George Dimitriadis, Joana Neto, Adam R. Kampff

## Abstract

Electrophysiology is entering the era of ‘Big Data’. Multiple probes, each with hundreds to thousands of individual electrodes, are now capable of simultaneously recording from many brain regions. The major challenge confronting these new technologies is transforming the raw data into physiologically meaningful signals, i.e. single unit spikes. Sorting the spike events of individual neurons from a spatiotemporally dense sampling of the extracellular electric field is a problem that has attracted much attention [22, 23], but is still far from solved. Current methods still rely on human input and thus become unfeasible as the size of the data sets grow exponentially.

Here we introduce the t-student stochastic neighbor embedding (t-sne) dimensionality reduction method [27] as a visualization tool in the spike sorting process. T-sne embeds the n-dimensional extracellular spikes (n = number of features by which each spike is decomposed) into a low (usually two) dimensional space. We show that such embeddings, even starting from different feature spaces, form obvious clusters of spikes that can be easily visualized and manually delineated with a high degree of precision. We propose that these clusters represent single units and test this assertion by applying our algorithm on labeled data sets both from hybrid [23] and paired juxtacellular/extracellular recordings [15]. We have released a graphical user interface (gui) written in python as a tool for the manual clustering of the t-sne embedded spikes and as a tool for an informed overview and fast manual curration of results from other clustering algorithms. Furthermore, the generated visualizations offer evidence in favor of the use of probes with higher density and smaller electrodes. They also graphically demonstrate the diverse nature of the sorting problem when spikes are recorded with different methods and arise from regions with different background spiking statistics.

## 1 Introduction

It is neuroscience dogma that the brain’s computational mechanics are implemented by the complex dynamics of its spiking neural networks. As a consequence, detailed knowledge of the spiking activity for ”as-many-neurons-as-possible” during behavior is seen as essential to understand how the brain receives and transforms information. Electrophysiological methods that record spiking activity extracellularly have been one of the most significant tools for exploring the correlations between behavior and neural activity and there has been a constant drive to record from more neurons, for longer times, from a host of neural regions, in diverse physiological conditions, and from many different species. This trend was recently accelerated by new microfabricated recording probes that extend the standard single electrode and tetrode devices [19] with integrated electronics to produce devices with thousands of recording sites [24, 1].

The new generation of recording tools brings with it the challenge of extracting meaningful physiological signals from the resulting (big) data sets. In the case of extracellular probe recordings, that usually means transforming the voltages measured at the electrode sites into spiking activity of the nearby neurons. The importance of accurate spike sorting stems from a number of ideas on how cell spiking contributes to brain functions. For example, competent sorting is required to test for sparse coding in memory function [6] or to assess the diverse responses of neighboring cells, important in theories of concept [21] and place cells [20].

The original attempts to spike sort greatly benefited from the development of the tetrode and its ability to simultaneously monitor the spiking signal of nearby neurons from multiple locations (i.e. 4) [10, 8, 29]. It has since become clear that dense electrode configurations, in which that same neuron is detected by multiple electrodes, generally improve sorting [13, 4], and hence the push for an increase in the electrode density of modern probe designs. Today new methodologies have evolved to work with the next generation of multi-electrode probes and to try and address the problem of the exploding size and complexity of the data sets [23, 22]. However, the basic idea of the spike sorting pipeline remains the same (Fig 1A). The (filtered) data go through a process of spike detection that has traditionally relied on thresholding the raw signal. The multi-unit activity generated is then passed through a dimensionality reduction method that transforms the space-time spike matrices into a smaller set of features. The most commonly used dimensionality reduction techniques are principal component analysis (PCA) [11] and wavelet decomposition [12, 18, 25] for offline and geometric/spike shape methods[9, 13] for online sorting. More recent approaches even combine the two offline methods to generate an optimum set of features for further analysis [21]. Finally, a clustering method is employed to automatically group together the spikes from an isolated single-unit in the high dimensional space of the decomposed features. Techniques commonly used for this clustering are k-means [30], mixtures of gaussians based on an expectation minimization algorithm [30] and template matching [33, 28]. An overview of different techniques for detection, feature extraction and classification is given in Bestel et al [3]. Methods that are currently under development follow a different route where the event detection, the feature extraction, and the clustering steps are realized in a single template matching step [16, 32]. These methods offer better parallelization capabilities and are proving very capable in handling millions of spikes arising from recordings of hundreds to thousands of channels. In all cases, the automated clustering algorithms operate on a number of dimensions that scales linearly with the number of channels of the recording probe. For the more recent multi-channel probes, this feature space usually contains hundreds of dimensions. Such multidimensional spaces make either manual clustering, or the manual supervision and quality assurance of the automated algorithms’ results, prohibitive. The t-sne dimensionality reduction technique was designed to reduce such multidimensional data sets to 2 or 3 dimensions in a way that visualizing them can offer meaningful insights into their original high dimensional structure [27]. Embedding techniques, like t-sne, transform the position of points in a high-dimensional space to positions in a lower (usually 2) dimensional space. This reduction transformation obviously requires that some information is lost. Each embedding technique decides which aspects of the original structure to keep and which to ignore. T-sne focuses on ensuring that the local structure (i.e. the ordering of distances between nearby points) remains intact while it ignores the global structure (i.e. the large distances in the t-sne space are not representative of the large distances in the original space). A good mental representation of how t-sne achieves this is to think of all points as objects connected to each other with spring-like forces. In the original space these forces are in equilibrium. When the points are transferred (randomly at first) into the 2D space the forces between them start both pulling and pushing so that a new equilibrium might be reached (Fig 1B). Points that are close in the original space are attracted to each other until they get roughly equally close in the 2D space, while points that are far away in the original space are repulsed by each other if they happen to find themselves close in the 2D space. This ability of the t-sne algorithm to repulse points that are nearby in the 2D space but not in the original space, offers a solution to the crowding problem of other embedding methods and underlies the informative 2D plots that it generates.

**Figure 1:**
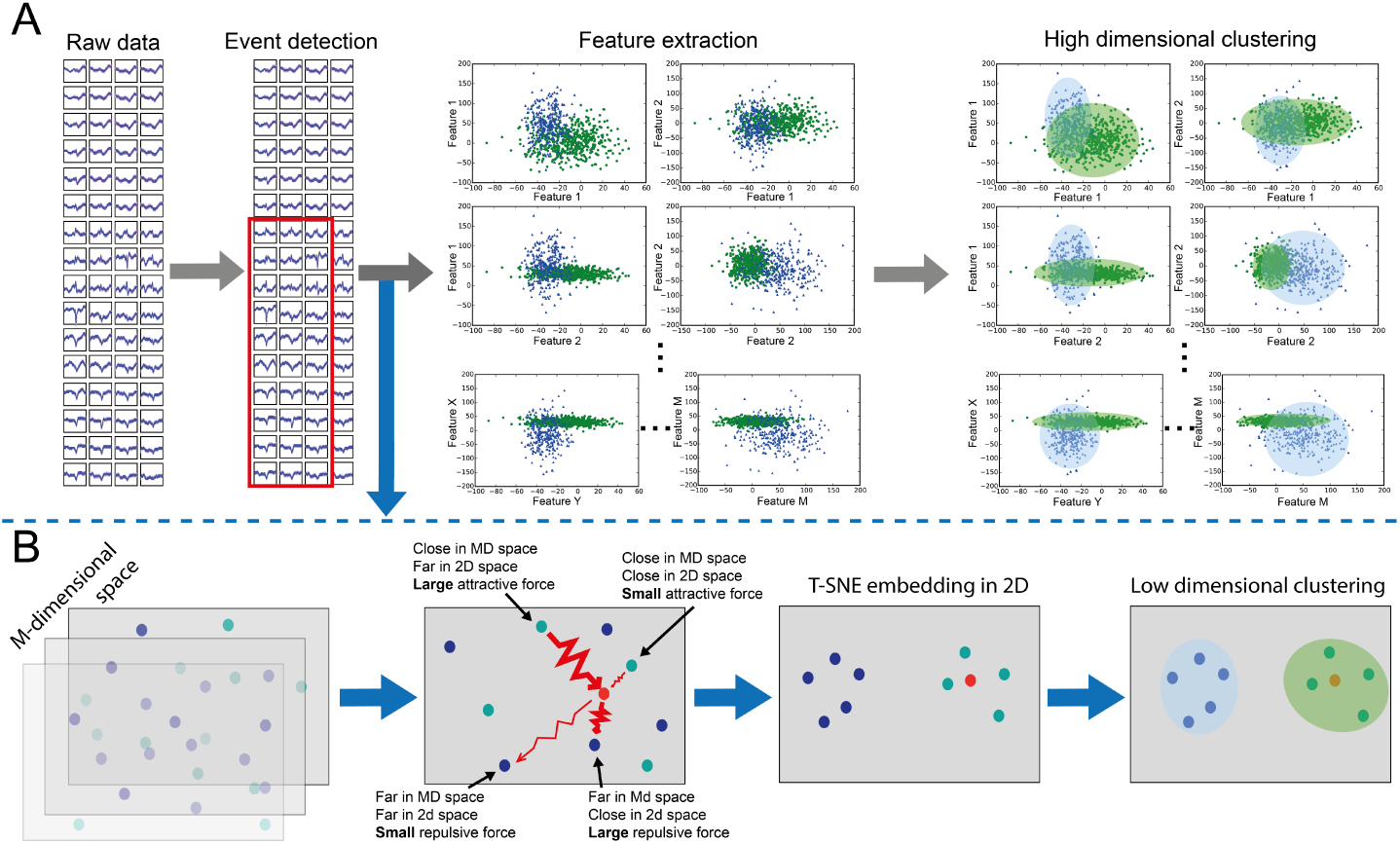
Graphical representation of A) the current pipeline for spike sorting and B) the way t-sne can be added as a visualization tool in this pipeline

In this work we apply t-sne to the spike sorting process and generate 2D plots that show obvious clusters of spikes. We use two types of data to validate our technique. The first is a ground-truth dataset that comes from paired recordings [15] with an extracellular and a juxtacellular probe, thus providing labels from the juxtacellularly recorded unit within the extracellular probe’s spiking activity. The second type is a hybrod dataset generated from the synthesis of real extracellular recorded data with manually superimposed spikes belonging to a number of single units [23]. In the following we demonstrate that many of the t-sne generated low-dimensional clusters represent the activity of single units, while others group together spikes arising from a large number of putative units and likely noise. We develop a GUI that allows the fast visual identification of the single unit clusters and report on how accurately the manually selected clusters represent the labeled single unit’s in our test datasets. We then use the visual representations of spike clusters that t-sne generates to offer an overview of how the sorting/clustering problem’s difficulty increases with decreasing electrode density. Finally, utilizing the input agnostic nature of t-sne, we use it to embed the results of a new template-matching algorithm (kilosort) applied to the same ground-truth dataset. We subsequently use the GUI to overview and manually correct kilosort’s results and show that t-sne’s 2D embedding visualization makes digesting and curating the high dimensional output of automated spike clustering algorithms a simple procedure and also provides a satisfying ”overview” of the otherwise overwhelmingly large, high-dimensional data sets. We will conclude with a discussion of possible extensions and future use cases of the t-sne algorithm for sorting and visualization of large-scale spike recordings.

## 2 Methods

### 2.1 Data sets

We used two types of data to test the efficacy of our methods. The first was a set of recordings from the anesthetized rat’s (motor) cortex [15]. In these datasets the extracellular probe’s recording was paired with a juxtacellular recording with a pipette. There were two different types of extracellular probes used in different sessions, a 32 channel staggered array (A1x32-Poly3-5mm-25s-177-CM32, NeuroNexus, USA) and a dense 128 channel matrix developed by the collaborative NeuroSeeker project (http://www.neuroseeker.eu). The 128 channel probe design is shown in Fig 4 and the 32 channel design in Fig S1. From the paired recording sessions available (www.kampff-lab.org/validating-electrodes) we selected one from the 32 channel probe (Paired Data 32: PD32) and one from the 128 channel probe (Paired Data 128: PD128). Those were the data sets in which the juxtacellularly recorded neuron was close enough to the extracellular probe to have its spikes easily detected on the extracellular recording without any spike triggered averaging. The PD128 set, when spikes were detected with 6.5 standard deviations threshold, consisted of 128820 spikes out of which 4420 where spikes belonging to the juxtacellularly recorded neuron (out of a total 4998 juxtacellularly recorded spikes). These are the spikes we used in the analysis shown in Fugures 2 and 4. At 4.5 std it consisted of 255026 spikes of which 4775 were the juxtacellularly recorded ones. When the same data set was put through the kilosort algorithm (where no detection through thresholding is actually used) it gave back 466313 putative ’spikes’ out of which we defined as non-spike noise the 139073 (see section 3.4), leaving 327240 spikes (72214 more than the 4.5 std threshold detection). The PD32 set consisted of 73814 spikes of which 331 belonged to the juxtacellularly recorded neuron.

**Figure 2:**
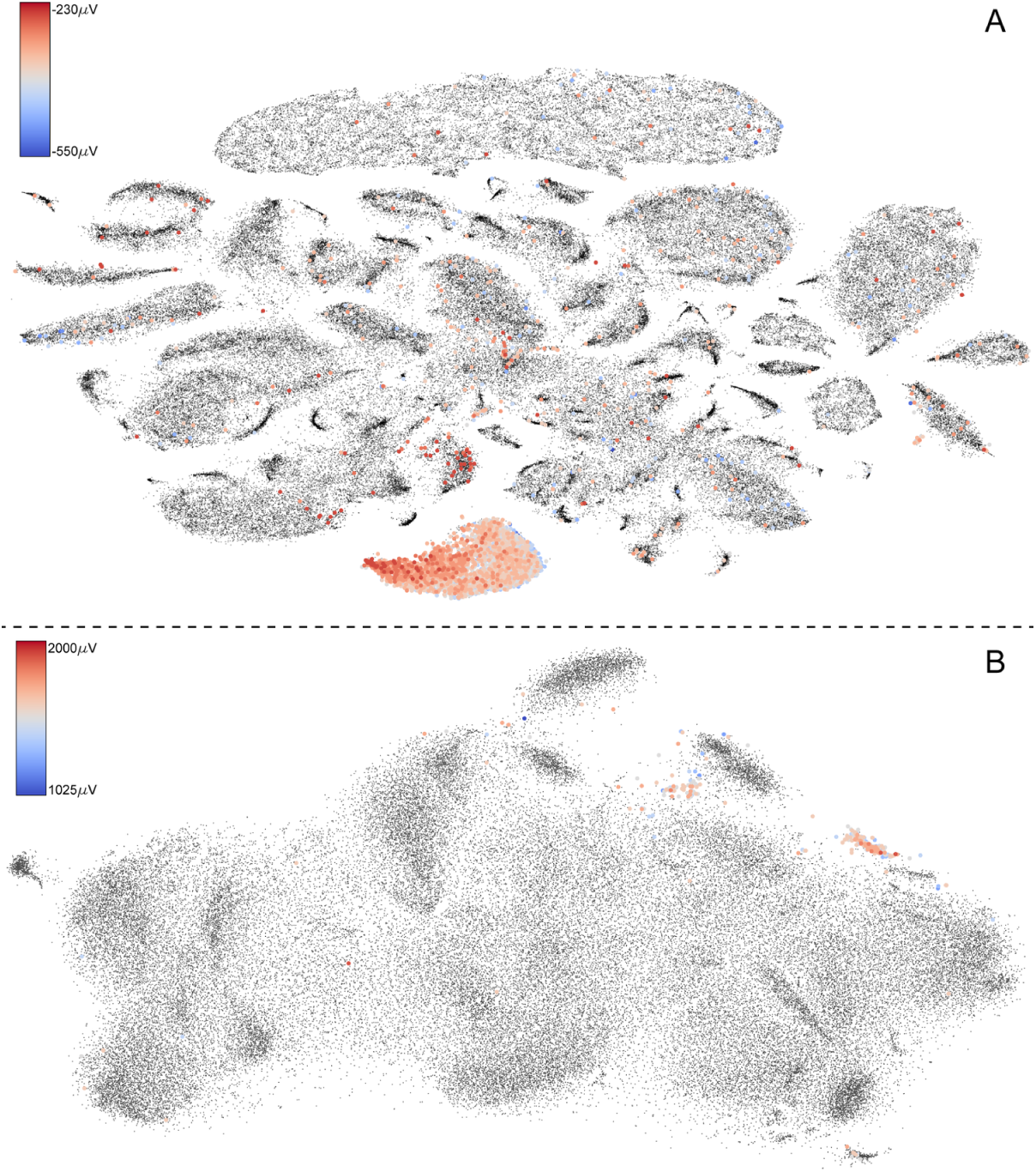
Results of t-sne analysis on PD128 (A) and PD32 (B). The color coding represents the size of the juxtaccellular spikes that correspond to the extracellular ones on the t-sne plots.

We used two hybrid data sets provided to us by the lab of Kenneth Harris. Their construction was based on recordings of an 129 channel probe and the superposition of previously sorted spikes derived from separate recordings (hybrid spikes). The details of how these hybrid time series were constructed can be reviewed in [23]. The first data set (HD1) had 86271 spikes with 7 hybrid spike sets coming from different (labeled) neurons ranging in size from 432 to 26043 spikes. The second data set (HD2) had 126102 spikes again with 7 groups of labeled hybrid spikes with group sizes between 442 and 24987.

### 2.2 T-sne code

For all our experiments we have used an extended version of the github available C++ t-sne code by van Maatens (https://github.com/lvdmaaten/bhtsne/). The bhtsne implementation in this repository allows for faster computations and makes larger data sets feasible through the use of the barnes-hut (bh) algorithm [26]. The bh algorithm groups together distant samples into a single average sample, minimizing the number of euclidean distance computations the algorithm must perform. This step makes the calculation of the actual error minimization of the 2D landscape faster and tractable for large sample numbers (on the order of millions). Yet the calculation of the initial probabilities in the multidimensional space still requires the algorithm to calculate the euclidean distances of all sample pairs adding a significant amount to the algorithm’s run time. To speed up this part we have implemented the computation of the euclidean distances between all sample pairs directly on the GPU. To do this we used the GPU euclidean distance implementation presented by Chang et al [5]. This allows us to run data sets of more than 10^5^ of spikes on a gaming desktop (i7 CPU, Titan X GPU, 64GB RAM) in just a couple of hours (see table ST1 about times for different number of samples and perplexity settings). However, storing the distances for all sample pairs on RAM soon imposes a bottleneck on the amount of samples that can be used. For N samples one requires 4*N*N (for 32 bit floats) bits of RAM, which translates to 40GB for 10^5^ spikes and to 1TB for 0.5*10^6^ samples. To overcome this bottleneck we have added the possibility to calculate the pairwise distances in groups and keep in memory only the ones that the barnes-hut algorithm will use (populating the tree structure used in the algorithm with GPU pre-calculated distances). This process requires a sorting step of the distances that increase the time required, but allows much larger sample numbers to be used.

For data sets that are prohibitively large, and to show that online sorting of samples is possible, we have extended the algorithm to be able to position samples on a t-sne pre-computed 2D landscape. This is done by measuring the euclidean distance of each new sample to the samples already passed through the t-sne algorithm. Then the extra samples’ positions on the 2D landscape are calculated as the average of the positions of the closest 5 original samples for each new sample. For spike sorting, given that the number of spikes passed through the t-sne algorithm offers a complete representation of the spiking units in the recording, we show that the algorithm correctly places the extra samples (see Supplementary Figure 2).

We have also extended the python wrapper that came with the original code to accommodate the use of the extra parameters our C++ t-sne function requires and to allow the user to choose between the use of the C++ executable or the CPU only scipy implementation of t-sne. The latter allows users without access to the CUDA library, or with hardware not capable of supporting our code, to still run a full t-sne spike sorting session only on CPU and fully in python (no C++ executable is called with this option), albeit for small number of spikes and with large run times.

### 2.3 Spike sorting pipeline

We have applied the t-sne algorithm in two separate points of the spike sorting pipeline using two different spike representation feature sets as inputs to the algorithm. In the first instance (results in Figures 2, 3 and 4) the algorithm operated on the masked PCA components of the spikes detected as described in [23]. In order to facilitate the input to t-sne of these components, we wrote python code that accepts the (masked or not) PCA components of the detected spikes, transforms these into a data set ready for the t-sne algorithm and calls the executable on it. We have also tried using raw data or non-masked PCA components as input to the t-sne algorithm, but found the masked-PCAs to consistently outperform these other inputs (results not shown). The second input feature set to the algorithm (results in Figure 5) was the distance to templates generated by the kilosort algorithm. In this case, the t-sne algorithm operated on feature vectors that measured how close or far each spike was to the set of spike templates that kilosort generated from the data.

We have also developed a minimal, and easily extensible, graphical user interface (GUI) that allows users to visualize the results of the t-sne algorithm and use this visualization to manually sort the detected spikes. The GUI offers a number of views and tools for manual spike sorting. These include: A 2D scatter plot of the t-sne results with several ways to select groups of spikes directly on the plot. A view to preview the selected group’s average time traces for all channels. The presented data are not explicitly filtered, but the baseline is subtracted using the first 10 samples. An autocorrelogram view of the selected spikes with 1 ms resolution bins. A heat map view of the average difference between the minimum and the maximum values in the spikes’ time window superimposed on the probe’s layout diagram. A way to label a selected group and store it as a cluster in a pandas structure (saved to disk as a pickle). The GUI also allows selecting and previewing of all plots for any previously saved cluster, deleting a selected cluster and previewing all saved clusters on the t-sne scatter plot. A number of input boxes and buttons also allow the merging and splitting of clusters as well as the reassignment of spikes to different clusters.

The development of the GUI is based on the bokeh python library. This makes further development of good quality views as desired by individual users relatively easy and fast. At the same time the bokeh library will struggle with large number of points presented simultaneously on the t-sne view. For the gaming desktop described above the easy to work with limit was reached at about 500,000 spikes.

The python code design was informed by the need to keep the code simple and extendable. To achieve this we chose to implement only functions, without any obfuscating object orientation or passing data around data structures in more complicated ways other than function arguments and return statements. Individual functions are constructed to be self-standing and usable outside the context of the spike sorting workflow the code was designed for. For example the basic t-sne functions can operate on any other data of a samples x features dimensionality, or the functions that produce the average spike timecourses or the average heatmaps of the probes can easily be used to generate plots outside the confines of the gui.

### 2.4 T-sne parameters and accuracy measurements

For spike detection we used a high and low detection threshold of 6.5 and 2 std respectively. We tried out a large range of t-sne parameters (perplexity, learning rate, theta and number of iterations) in order to define the set that gave us good results, but that didn’t take too long to run. For all of the results presented here, the parameters used were perplexity 100, learning rate 200 and theta 0.2. The theta parameter defines the angle of the cone inside which all points are treated as a single average point by the bh algorithm. Smaller values mean that the algorithm averages fewer points together, i.e. only those that are far away from the central point. A value of 0.2 is considered an approximation closer to an exact solution. For the PD128 and the HD1 sets we ran the algorithm for 2000 iterations while for the PD32 and the HD2 for 5000. Perplexities lower than 100 were shown to compromise the results (in as far as separation of clusters defined by visual inspection) while higher numbers (we tried up to 1000) would make no obvious difference other than adding to the run time of the algorithm. We have found that perplexity is a sample number dependent measure, but that for tens to hundreds of thousands of samples (as in all our data sets) the chosen number of 100 offered the best quality vs run time balance. The fact that over a certain value perplexity did not seem to change the quality of the embedding, adds to the idea in the t-sne literature that this is a stable parameter that can vary a lot without substantially influencing the results.

Having labeled data allowed us to measure the quality of the t-sne clustering visualization as a tool for separating single units. Since the t-sne algorithm itself does not cluster (i.e. label) the data, but only offers a 2D embedding, we needed a way to label the spikes according to their position in that embedding. We chose to use the density-based spatial clustering of applications with noise [7] (DBSCAN) algorithm since it provides a non-parametric way to label the embedded spikes by clustering together samples that form denser groupings compared to their immediate environment. We found DBSCAN’s approach to clustering matching most closely the human intuition of neural units corresponding to separate groups of spikes in the 2D visualization of the t-sne data.

Having established a method for labeling the t-sne results we then compared the generated labels with the ground truth information from the juxtacellular recordings or the hybrid spike groups. We report here the results of three commonly used measures for such comparisons. The first is Precission (or Confidence or True Positive Accuracy) being the ratio of the true positive samples (i.e. spikes labeled by DBSCAN as part of a unit that also had either a juxtacellular spike correspondence or the correct hybrid label) over all positively labeled samples (all spikes defined by DBSCAN to belong to the specific single unit). The second is Recall (or Sensitivity or True Positive Rate) being the ratio of the true positive samples over all true samples (all spikes with a juxtacellular spike correspondence or a specific hybrid spike label). The third is the F-factor which is defined as the harmonic mean of Precission and Recall (i.e. 2*Precission*Recall/(Precission+Recall)). We also calculated the Receiver Operating Characteristics (ROC) values for each label (either hybrid spike set or juxtacellular corresponding set) as a point on the plot of the True Positive Rate versus the False Positive Rate (see Supplementary Figure 2). The False Positive Rate is defined as the ratio of the false positive samples (spikes defined by the DBSCAN as part of the label but not having a corresponding juxtacellular spike or a hybrid set label) over all the negative samples (all spikes not having the specific juxtacellular or hybrid label).

## 3 Results

### 3.1 Paired Data

We ran the t-sne algorithm (for a 2D embedding) on the full 128820 spikes of the PD128 set; each spike was represented as a masked vector of 384 dimensions (128 channels * 3 largest PCA components per channel). The masking of the PCA components (i.e. modulated between 0 and their full value) is described in Rossant et al. [23]. The resultant embedding is shown in Figure 2A. The embedding generates a number of distinct groups of spikes and a number of more diffuse clouds with varying internal densities. A spike grouping at the bottom of the figure contains the majority of the extracellular spikes that correspond to juxtacellularly recorded spikes (i.e ground truth spikes from the isolated single cell). The figure shows that a small percentage of the spikes coming from the labeled cell are spread throughout the entire 2D space, while the labeled cluster also contains some spikes that are not generated by the labeled cell. For this set of ground truth spikes the Precission, Recall and F-factor are 0.86, 0.83 and 0.84 respectively, which translates to 14% of the ground truth spikes not being classified as part of the main cluster and to 17% of the main cluster spikes being misclassified as part of the ground-truth.

The color coding of the samples with a corresponding juxtacellular spike shows that the embedded cluster’s internal structure has a strong relationship with physical characteristics of the actual spikes, in this case their peak amplitude. The t-sne algorithm is designed to retain the local structure of the multidimensional space during its transformation into the 2D embedding space. That means that any correspondence between the physical properties of the samples and their distances in the high dimensional space of their PCA components will be retained in the subsequent embedding, at least for the samples that are close together (i.e. the ones within a single cluster). The 2D embedding space is easy to visualize and thus allows quickly noticing such relationships. By simply changing the color coding scheme to represent different properties one can easily browse the relationships (or lack thereof) between any number of specific characteristics. For example, color coding the spikes according to the time they appear in the recording reveals that there is no correspondence between the size of spikes and the time they were recorded (results not shown).

The clearest picture of how t-sne operates on the data can be gained from videos of the t-sne process in which each frame is the result of progressive iterations of the embedding algorithm. We captured this process for the PD128 data set in Supplementary Video 1.

Figure 2B shows the t-sne embedding of the 73814 spikes of the PD32 set. Here the embedding generates a more homogeneous cloud with significantly fewer easy to delineate groupings of spikes. The Precission, Recall and F-factor for the juxtacellularly labeled spikes in this case are 0.91, 0.65 and 0.76 respectively for the t-sne/DBSCAN generated cluster with the most color coded spikes. In this case the cluster is a fairly homogeneous one (with only 9% of the spikes not belonging to the unit it represents) but it fails to capture a large percentage (%35) of spikes from the same unit which end up in other groups. Also, the cluster’s internal structure shows no correspondence with the spikes’ peak amplitude as measured by the juxtacellular electrode. We will propose (see Probe electrode densities and clustering quality) that the drop in clustering performance between the P128 and P32 embeddings is mainly due to the lower sampling density of the extracellular space provided by the 32 channel probe.

### 3.2 Hybrid Data

We also used t-sen to embed the two hybrid data sets described in Methods / Data Sets. The results for HD1 can be seen in Figure 3A and for HD2 in 3B. In this case, the output of the t-sne algorithm provides a clean visualization of the known single unit clusters. Most clusters are fully separated from the other clusters and contain a very small number of spikes that do not belong to their corresponding unit. The spikes shown in black in both sets are not hybrid spikes, but rather those that existed in the original recording, i.e. not ground-truth. For HD1 all clusters have Precision, Recall and F-factor of 0.99. For the HD2 set, clusters 1, 4, 5 and 7 have Precisions, Recalls and F-factors ranging between 0.95 and 0.99. Cluster 6 has 0.91, 0.9 and 0.91 respectively.

In the case of cluster 5, DBSCAN did not classify as part of the unit the spikes that expand away from the main group to the right of the cluster. Yet their proportion was small enough to keep the Recall at 0.95. The same applied to cluster 1, which was missing the group of spikes showing at the top of the large not Ground Truth spike grouping (black group at the left of the plot labeled as Not GT). That group of spikes was also small enough in percentage that the cluster still achieved a Recall of 0.96. T-sne failed to fully separate clusters 2 and 3 (see insert of Figure 3B).

**Figure 3:**
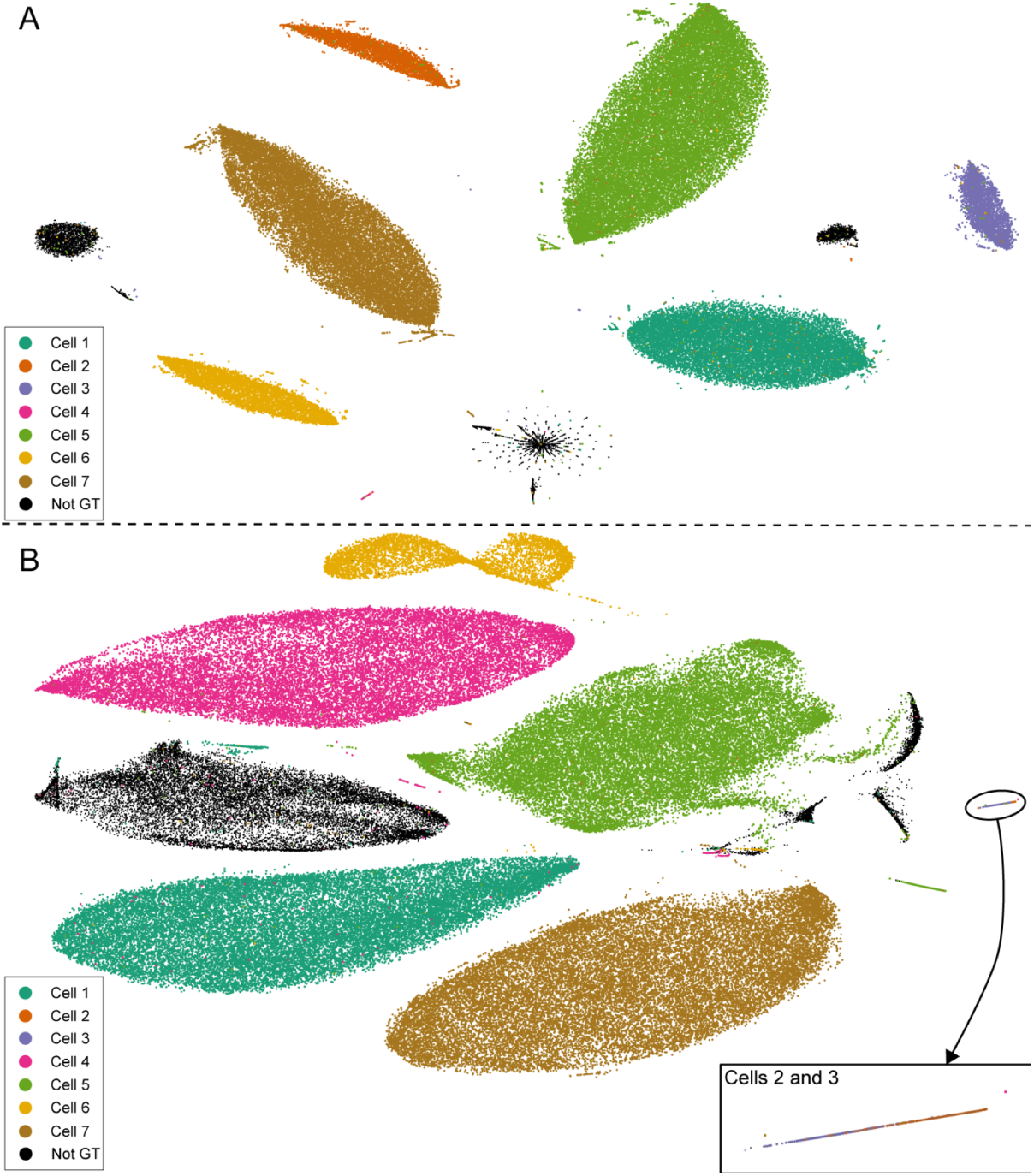
Results of the t-sne analysis on the hybrid data, HD1 (A) and HD2 (B). The colors represent spikes that belong to pre-classified units added to the data sets while the black points represent spikes pre-existing in the datasets and thus do not belong to known units. The insert in B) shows a zoom-in of the embedding of units 2 and 3 that t-sne does not manage to segregate.

In this case the clusters’ Recalls were 0.99 but their Precissions were 0.44 and 0.55 with F-factors of 0.61 and 0.71 respectively. The internal structure of their common spike group in the 2D embedding space shows that the similarity of their spikes changes in a uniform way and that there are some spikes from the two groups that are quite similar, thus causing the embedding to overlap the two units.

### 3.3 Probe electrode density and clustering quality

As the number of electrodes per probe increases, it is important to understand how the density of electrodes relates to an algorithm’s ability to reliably extract and isolate single-units. A major argument for the miniaturization of silicon probes was the idea that more densely spaced electrodes will increase spike sorting quality by providing more features with which to classify spikes. Here we review this argument using t-sne by performing embeddings of the P128 dataset, originally captured with a density of 2050 electrodes/mm^2^, at artificially reduced densities. This was achieved by removing more and more channels while keeping the total coverage intact and then performing a new t-sne embedding with each sub-dataset. The results visually demonstrate how artificially decreasing the probe’s density affects the sorting of the detected spikes.

We started from the PD128 set and used the manual sorting GUI we developed (see Methods) to define as many groups of spikes belonging to the same unit. The criteria used to delineate a number of spikes as a single unit were the following: The spikes had to be close in the t-sne space, ideally belonging to one, obvious, unique grouping (e.g. large orange cluster at the right of the plot or yellow cluster at the bottom), but if not so, the grouping had to at least be contiguous (e.g. the dark red and green clusters at the far right and middle of the plot or the yellow and red clusters at the left and top of the plot). The average time course of the spike on all electrodes had to show a standard extracellularly measured action potential. The autocorrelogram of the spike times within a group had to be zero within 1 ms od time zero (i.e. no two spikes of the group could have been fired in a time interval sorter than 2 ms of one another). Finally, the heatmap of the group had to show a contiguous region of activation on the probe that was compact (just a few neighboring electrodes). Following the above criteria we labeled 90 units in the PD128. Given the extent and density of electrodes as well as the position of the probe within cortex, this is consistent with the number of neurons one might expect to detect in local vicinity, i.e. ¡ 50 microns [15]. Figure 4A overlays the manually classified clusters directly on the t-sne plot. There are three separate groupings of spikes that we have not labeled as a single unit (the grouping at the top of the plot and two on the right side of the plot between the orange, yellow, blue and green units). T-sne in this case had clustered together three groupings of multi unit spikes. These groupings showed also no internal grouping that could be assigned to a single unit.

Figure 4B to 4F show the t-sne results arising from the same dataset with some electrodes removed. From B to F the number of electrodes used were 64, 32, 22, 16, and 8 respectively (denoted as blue in the figure). The color coding in these subplots is the same as in Figure 4a and denotes the unit each spike belongs to as defined by the above procedure. Even at 64 channels there is an obvious deterioration of the t-sne clustering quality that progresses all the way to the 8 channel data set. This deterioration can be seen as a reduction of the number of easy to delineate spike groups, an increase of the number of spikes that form part of the larger amorphous clouds, and by the mixing of the labeled spikes amongst the unit clusters and between the clusters and the undifferentiated cloud structures.

**Figure 4:**
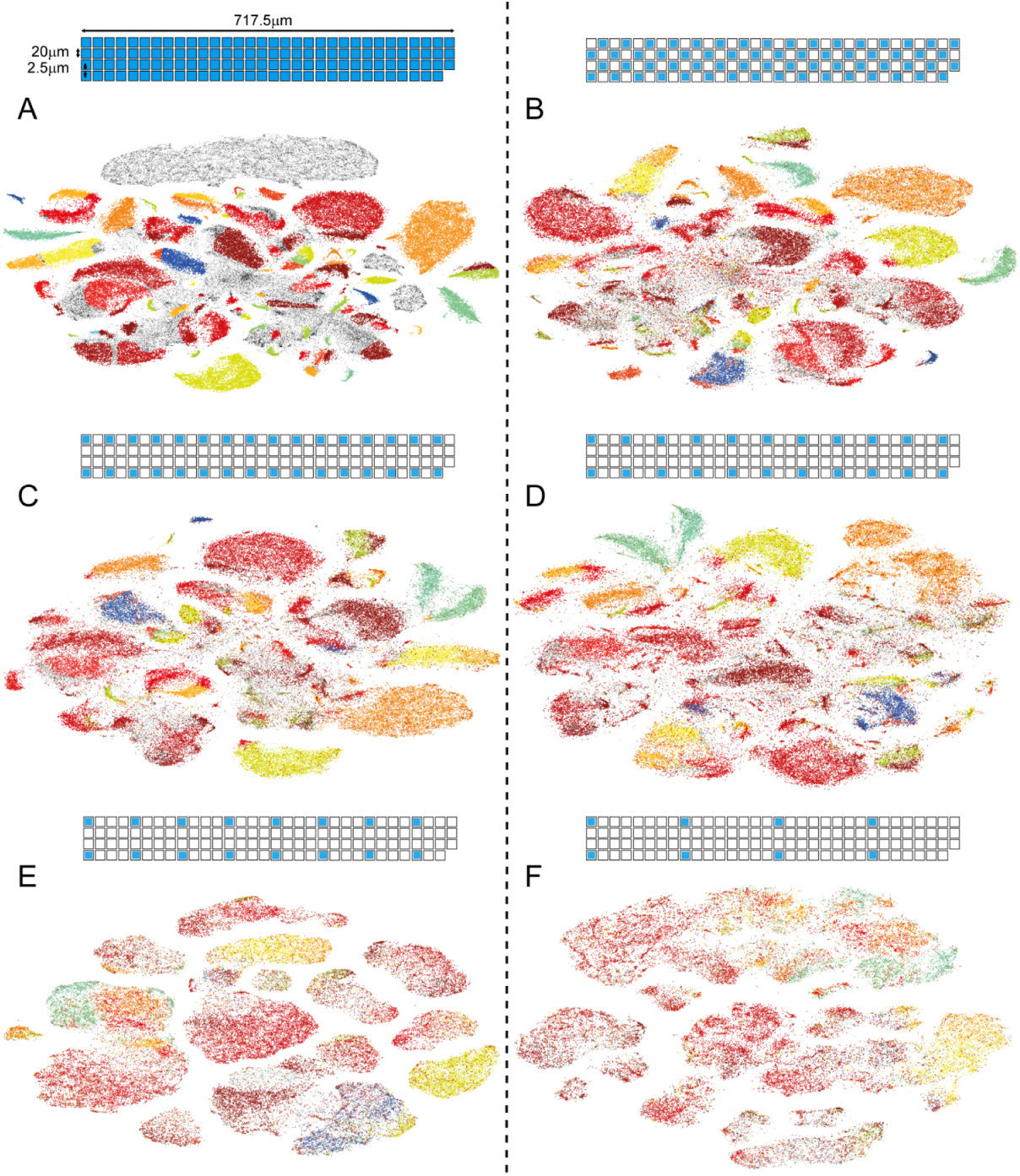
T-sne embeddings of the same PD128 data set for a series of electrode densities (the electrode configuration is shown at the top of each subplot with blue being the used channels). The color code represents the manually sorting of the spikes done on the full (128 channels) t-sne embedded data set. The way the different color groups start to break up and merge with each other over the consecutive electrode configurations visually depicts the increased difficulty of the spike sorting problem for probes with reduced densities. For the last two densities (subplots E and F) the electrodes are far enough from each other to ensure that each spike is seen by only one electrode. The quality of these embeddings makes immediately obvious the usefulness of the multiple, spatially separated, channels with information about each spike.

### 3.4 Intuitive manual curation of automated clustering

Template matching algorithms are proving to be fast alternatives to the more classical detect, embed, and classify spike sorting pipeline, especially for probes with a large number of densely spaced electrodes. These algorithms iteratively generate templates of the spikes present and then use these templates to detect and classify the individual spikes. The result is a list of spike templates and the euclidean distances of each spike to each of these templates. Each spike is assigned to the template that matches it the best (i.e. it has the smallest distance to). We report here how t-sne embeds these template distance features and how we utilized our GUI to sanity check the automatically generated results and get a broader overview of the clustering problem for a specific data set.

We ran the kilosort algorithm [16] on the PD128 data set with a maximum number of templates set to 256 (double the number of channels on our probe). It generated 252 templates and detected 466313 spikes. We then used the distances of each spike to all the templates as the t-sne input feature space (a 252 dimensional one), setting to 0 all the distances except the nearest 16 templates. The result of this t-sne embedding can be seen in Figure 5. The embedding shows three different types of spike groupings. The first are a number of very tight groupings (denoted from here on as ”points”) where the spread of the entire group in both the x and y axis is at least 3 orders of magnitude smaller than the total spread of the embedding. The second is a linear grouping where there is an obvious extended axis to the grouping (not necessary along the x and y axis of the full embedding), one with a very small and one with a much larger spread (denoted from here on as ”lines”). The third type of grouping has large spreads on both the x and y axis, no less than 2 orders of magnitudes smaller than the total t-sne spread (denoted here on as ”blobs”).

**Figure 5:**
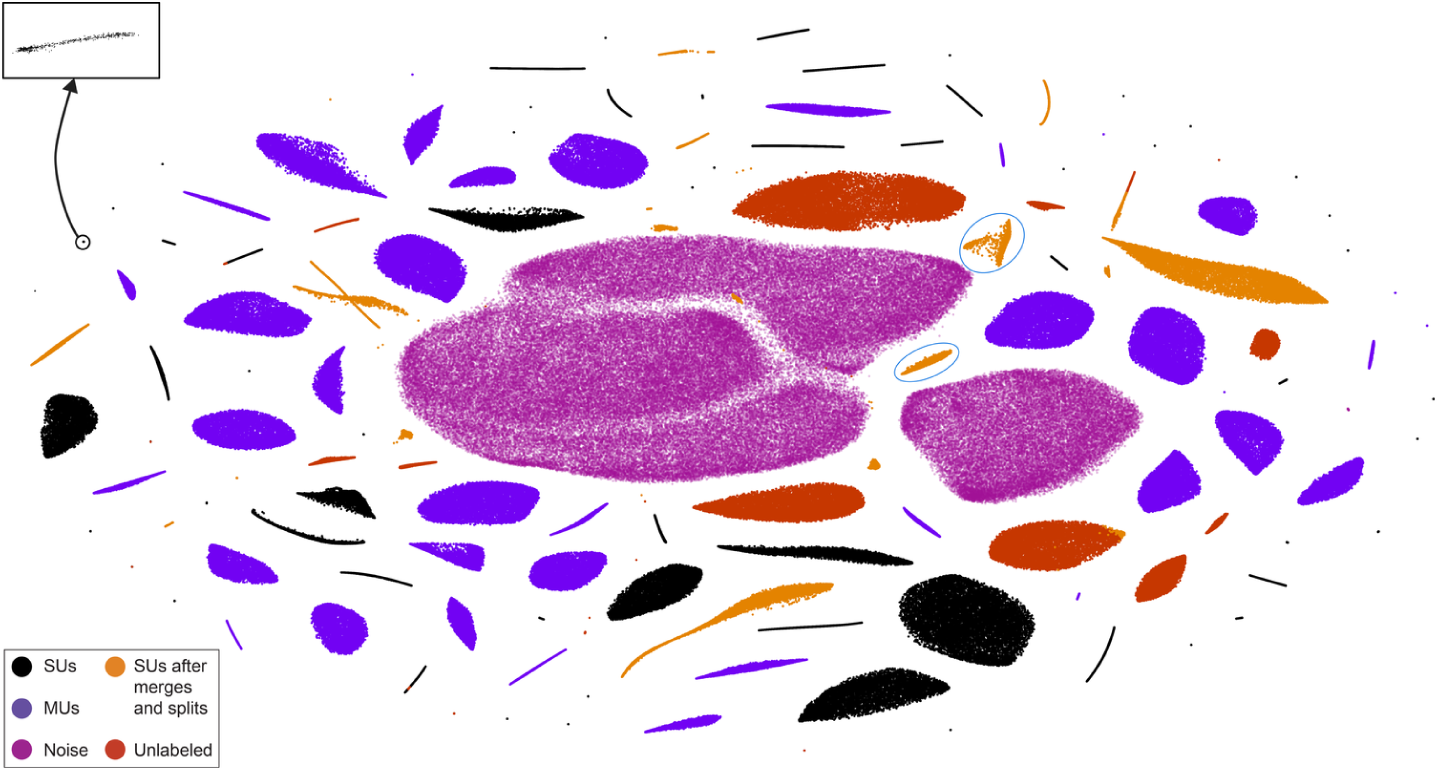
T-sne embedding of the results of the kilosort algorithm (distances to templates) for the PD128 data set. Colors indicate the different categories that the spikes fall into after using our custom GUI to manually evaluate and correct as necessary kilosort’s spike assignments to templates. The SU labeled groupings represent a single unit each. The MU labeled groupings represent a single template each but multiple units. The Noise labeled groupings represent noise that has been picked up by kilosort as spike templates. The SUs after merges and splits have a more complex representation where each single unit can be either one or more groupings and have originally belonged to one or more templates. The Unlabeled groupings represent spikes that could not be assigned to either a single unit, or a multi unit or a noise group. The insert at the top of the figure depicts a zoom-in (x30 magnification) to a point grouping (SU) showing that these groupings are actually comprised of large number of spikes all embedded very close to one another. That compactness indicates a very large similarity in the template distance feature space and a large dissimilarity to any other spike, so it is not surprising that they are the groupings that kilosort has assigned to templates that represent single, easy to define units.

Using the manual clustering GUI we could easily evaluate in which kilosort assigned templates the spikes in the different embedding groupings belonged and vise versa (i.e. which groupings held spikes from any single template). From this comparison five general categories emerged. The first category involved groupings that was fully represented by a single template and whose heatmap and autocorrelogram both indicated a single unit. There were 73 of these single units (SUs) out of which the majority (49) were point groupings, 18 were line groupings and a small minority (7) were blobs. This category included 109890 spikes. The second category were groupings that again was fully defined by a single template but this time the autocorrelogram (and sometimes the heatmap) indicated multiple units (MUs). There were 39 of these MUs, the majority of which (26) were blob groupings with only a small minority being either lines (7) or points (6). There were a total of 134288 spikes in this category of MUs. The third category was represented by the three largest, semi-connected, blob groupings in the center of the embedding and the 4th largest blob on their right. Their ’spikes’ belonged to 5 templates (4 for the central blob and one for the blob on the side) and each template had its ’spikes’ grouped together but with a large minority spread throughout other parts of the groupings. These 5 templates were the most numerous and they all showed average spike trains that were very small in amplitude (¡ 100uV), did not resemble spike shapes, and were identical on all channels. We denoted this category as noise. It included a total of 139073 ’spikes’.

In the fourth category we lambed together all the cases that after splitting or merging of templates or moving spikes from one template to another ended up with acceptable SUs (based on their average spike shape, their probe heatmap, and their autocorrelogram). We arrived at 24 SUs that included a total of 38482 spikes. In these cases the combined information of where the spike lay in the embedding and in which template it belonged to made it fairly straightforward to either appropriately merge or split templates or move spikes along templates. In some cases a single SU would be represented by two separate t-sne groupings. An example of this are the two groupings embedded within blue ellipses in Figure 5. They represent a single unit after the merging of two kilosort templates (with spikes mixed over the two groupings). This unit happens to correspond to the juxtacellularly marked spikes. The unit has 4845 spikes in it. Out of those, the 4821 spikes’ timestamps correspond to the 4998 juxtacellular timestamps within a jitter of 1 ms. That translates to a False Possitive error of 0.5% and a False Negative error of 3.7%. In other cases, a single grouping would contain spikes assigned to two templates and a merging indicated a single SU (like the orange line at the bottom center of the embedding) while in others a single template would be represented by multiple groupings each defining an SU, resulting in the template’s splitting (like the two orange line groupings at the top center of the embedding). Finally the 5th category involved all the spikes that we were unable to assign to any of the previous divisions (SUs, MUs or noise). These were spikes that showed no obvious correlation between the embedding position and their template assignment. For example, the large red blob to the top and right of the large noise blob had 14184 spikes that kilosort had assigned to 95 separate templates (most of which were templates with fewer than 10 spikes each). There was no internal structure to the blob, i.e. the spikes of each template appeared randomly spread throughout. The embedding of all these spikes and templates in a single grouping made it straight forward to visuallize the situation and assign all the spikes to the unlabeld division.

## 4 Discussion

Spike sorting has evolved over the last 20 years from a set of techniques to discriminate 5 to 10 distinct units in a space of a few tens of features to strategies for labeling many tens to hundreds of units in large feature spaces. However, the number of distinct units is expected to soon reach well into the thousands. The emerging demands these growing datasets have inspired the development of more capable automatic sorting algorithms. However, the manual overview of the spike sorting process remains an essential step of the pipeline, albeit one that is becoming increasingly labor intensive and error prone. The problem of producing human readable visualizations of structures that exist in large dimensional spaces is of course not unique to spike sorting but is common in all data intensive fields. One commonly employed strategy for working with such data is the use of non-linear dimensionality reduction methods that try to retain in their projections as much of the initial structure of the data as possible. The current state of the art in these methods is t-sne, which manages to project onto two or three dimensions the multi-dimensional data in a way that preserves its local structure and makes the visualization of this structure both possible and intuitive. A large literature has evolved applying this technique to a diverse number of big data problems ranging from AI [14] to genetics [31, 17], and behavior [2]. We show here that, in the case of spike sorting, using t-sne on the results of either a masked PCA or of template matching, provides a visualization that offers a clear picture of how the individual spikes form groups in the high dimensional feature space. These groups directly relate to the individual units that generate the spikes, which can now be visualized and curated in an intuitive manner or compared to the results of an automatic clustering method. Furthermore, t-sne allows a visual inspection of the structural groups that different data sets of spikes contain, something that can be achieved neither by looking at 2D plots of pairs of features within a multi-dimensional space or by any current clustering algorithm. Here we demonstrate that t-sne visualization immediately reveals differences between biological and hybrid datasets (Figure 3), how spike sorting challenges accrue when each spike is represented by features generated by fewer and fewer electrodes (Figure 4), and that manual curation of the output from an automated template matching procedure becomes effortless in a low dimensional space (Figure 5).

The generality of the t-sne method also makes it applicable to datasets that arise from diverse sources. The successful application of t-sne to behavioral data [2] suggests the possibility of applying it to datasets comprised of a combination of spike and other electrophysiological signals (LFP, etc.) as well as the behavioral features occuring during the recorded neural activity. It is thus possible that groupings in the t-sne visualization of such a dataset could provide informative clues as to the connection between different forms of brain activity and to behavior itself.

Finally, we are now working on using t-sne embeddings into three-dimensional space to generate spatial (X,Y,Z) representations that can be explored in virtual reality, thus providing a new form of access to complex datasets. (Supplementary Video 2)

## 5 Additional information

### 5.1 Software access

A first version of the software used here can be downloaded through the following means. In a conda environment do: ”conda install -c georgedimitriadis t_sne_bhcuda”. That will install both the python code and the cuda executable and will work in either Windows or Linux OSes. You can grab the python code from the pypi repository:

https://pypi.python.org/pypi/t_sne_bhcuda/0.2.1.

You can get the C / CUDA code for the cuda executable from the github repository:

https://github.com/georgedimitriadis/t_sne_bhcuda.

We are currently in the process of re-writing the code so that the cuda part is embedded in the python code. For this and future developments check the pypi and github repositories or contact the authors.

